# Reaching to inhibit a prepotent response: a wearable 3-axis accelerometer kinematic analysis

**DOI:** 10.1101/2021.01.28.428591

**Authors:** Alessia Angeli, Irene Valori, Teresa Farroni, Gustavo Marfia

## Abstract

The present work explores the distinctive contribution of motor planning and control to human reaching movements. In particular, the movements were triggered by the selection of a prepotent response (Dominant) or, instead, by the inhibition of the prepotent response, that required the selection of an alternative one (Non-dominant). To this aim, we adapted a *Go/No-Go* task to investigate both the dominant and non-dominant movements of a cohort of 19 adults, utilizing kinematic measures to discriminate between the planning and control components of the two actions. To sample such measures, a low-cost, easy to use, 3-axis wrist worn accelerometer was put to good use to obtain raw acceleration data and to compute and break down its velocity components. The values obtained with such task indicate that with the inhibition of a prepotent response, the selection and execution of the alternative one yields both a longer reaction time and movement duration. Moreover, the peak velocity occurred later in time with respect to the dominant response, revealing that participants tended to indulge more in motor planning rather than in adjusting their movement along the way. Finally, comparing such results to the findings obtained by other means in literature, we discuss the feasibility of an accelerometer-based analysis to disentangle distinctive cognitive mechanisms of human movements.

## Introduction

Our everyday life is deeply defined by the voluntary actions we execute towards ourselves and towards the world that surrounds us. The way we plan and control our movements has been widely investigated for different motor tasks, to deepen our understanding of which motor strategies individuals adopt to select and execute different goal-oriented actions. In particular, as action features are usually movement-specific, this work focuses on a specific arm movement, namely reaching, which allows human beings to act within their peri-personal space by grasping, manipulating and using objects, as well as to interact with their own bodies and with other people.

Performing this specific action requires both a pre-planning and an on-line control of the desired motor output. Such two mechanisms are settled in distinct brain regions, respectively intervene in either the early or later movement time and appear influenced by different sensorimotor features and cognitive processes of the action [1]. Indeed, the role of motor networks might go beyond the action specification that answers to the “how to do it” and contribute to the simultaneous process of action selection, which addresses the “what to do” issue and chooses among currently available options [2]. It goes without saying that cognitive control is fundamental to the process of action selection, including the ability to inhibit inappropriate or incorrect responses [3]. For this reason, researchers look forward a further comprehension of how people perform or inhibit reaching movements.

In neuro-psychology, one of the most commonly used task to assess inhibition is the *Go/No-Go* paradigm [4]. The “Go” trials require participants to provide a response (i.e., do something) as soon as a dominant cue appears. On the other hand, the “No-Go” trials require to inhibit the response and not answer (i.e., do nothing) when another non-dominant cue appears (the latter usually appears less frequently than the dominant one). Reaction time (computed as the time elapsed from the cue onset to the response) and errors commonly characterize the performance of inhibition.

However, such a task is unable to investigate the different motor strategies individuals may adopt to perform either a prepotent or an alternative response. Therefore, the kinematic analysis of a reaching movement to select or inhibit a prepotent response would convey additional information on how it was executed and distinguish between planning and control aspects [5]. To this end, kinematic measures have been included with adapted *Go/No-Go* paradigms that asked participants to perform either a prepotent action elicited most of the time (dominant), or an alternative less frequent one (non-dominant). In one adaptation of the task, it was possible to analyse Reaction Time (RT) and Movement Duration (MD), which shed light on the distinctive motor strategies implemented for the prepotent or alternative response [6]. For instance, the non-dominant action might be performed with a longer RT or a longer MD depending on whether the actor required either a longer planning phase before the movement onset or a greater control and adjustment during its execution. Nevertheless, it is worth noting that motor planning is not relegate to RT but also overlaps with motor control during the MD. Indeed, “as planning is generally operative early and control late in a movement, the influence of each will rise and fall as the movement unfolds” [1, p. 5]. Therefore, kinematic indices other than RT and MD would be more informative to further investigate the way in which planning and control affect the movement execution. The following Section will expand on some kinematic parameters that can clarify the mechanisms beneath distinct movements. Such an approach is also promising to distinguish the specific impairments of inhibition skills, which are common of several neuro-psychological conditions [7].

Although motor analysis is highly informative both in research and clinical settings, kinematic studies often rely on expensive, bulky and sophisticated motion capture systems which may not be affordable in most operative and experimental contexts. In order to use low-cost portable solutions and boost the applicability of motion analysis, both custom made [8] and commercial tools have been recently evaluated. One extensively used commercial option is the Leap Motion Controller system, a small compact device containing two cameras and three infrared light diodes which has, however, spatial and temporal limits compared to motion capture systems [9]. Another commercial possibility that seems more promising in terms of measurement reliability and validity are the inertial sensors built with 3-axis accelerometers, gyroscopes, and magnetometers [10]. In light of these encouraging evidences, the time is ripe for the use of low-cost accelerometers to investigate the distinct neuropsychological mechanisms beneath action selection.

## Kinematic measures and related work

The movement research field has extensively debated regarding the distinctive meaning, reliability and validity of different motor indices. The total response time represents the time spent from the onset of a “Go” cue to the end of the movement. A preliminary distinction within the total response time individuates the RT and the MD [11]. RT includes the initial still phase that is devoted to detect and perceive the cue and also plan the appropriate movement towards the target. MD involves the movement execution, namely the time during which the actor moves to reach a target stimulus (either spontaneously or after a “Go” cue is shown). In addition, it is possible to find other indices depending on kinematic quantities, such as acceleration (e.g., mean and maximum acceleration, movements units - number of acceleration/deceleration phases, jerk), velocity (e.g., mean velocity, value of peak velocity, time to peak velocity) and displacement. For the purpose of investigating human movements, such indices are affected by different factors, thus providing insights regarding distinct neuro-psychological mechanisms underlying motor activities. For example, the motor or social goal of an action differently affects its deceleration time and peak velocity [12].

Acceleration, in particular, discloses the smoothness of a movement. The optimal reaching movement is ideally (for instance in experimental contexts and robotics) the one that maximizes the smoothness considering the jerk, namely, the rate of change of acceleration [13, 14]. The smoothness of a reach-to-grasp movement might depend on whether the target object is present, imagined or absent, on how it is oriented, or on which is the plane of movement (e.g., horizontal or vertical plane) [15].

Neuro-imaging studies examined the association between kinematic quantities and cortical activities and collected evidence of distinct cortical networks being related to distinct kinematic indices. This association has been investigated by Bourguignon et al. with the cortico-kinematic coherence analysis, which employed time-locked magneto-encephalographic (MEG) and accelerometer signals of movements [16]. For instance, fast repetitive voluntary hand movements of neuro-typical adults revealed that movement acceleration was mainly coupled with a coherent activation of contralateral primary motor (M1) hand area at ≈ 3 Hz and ≈ 6 Hz of movement frequencies. Moreover, only when the hand movement aimed at touching its own fingers, the primary somato-sensory (S1) hand area became the most coherent brain area at ≈ 3 Hz of motion frequency. In addition, also the activation of the DLPFC (dorsolateral prefrontal cortex, which is responsible for goal-directed action planning) and the PPC (posterior parietal cortex, which is responsible for sensorimotor integration and movement monitoring) areas were coherent with movement acceleration [17].

Focusing on velocity, the minimum-jerk model predicts that reaching trajectories starting and ending at full rest will show a symmetric, bell-shaped velocity path, with 50% of MD spent both accelerating and decelerating. However, MD and velocity across time are also shaped by several factors, such as the individual developmental trajectory [18], the affordances of the target object (e.g., a cup or a spoon) [15], and social intentions during interactions with others [19]. One of the indices that kinematic research uses to detect the effect of those factors is the Time to Peak Velocity percentage (TPV%). Given that whether a kinematic parameter occurs earlier or later over the MD would reflect more either planning or control [1], a small TPV% resulting in a longer deceleration phase may indicate a greater need for control and adjustment of the ongoing movement. On the other hand, a big TPV% resulting in a shorter deceleration phase may indicate a greater need for motor planning.

The aim of this project has been to evaluate the kinematics of reaching movements during a task that required neuro-typical adults to either select or inhibit a prepotent response. Kinematic indices such as RT, MD and TPV% were calculated to disentangle the contribution of motor planning and control in the selection or inhibition of a prepotent response. To the end of using a low-cost portable tool, we employed a 3-axis wrist worn accelerometer. In the following, the methods and results, as well as considerations concerning the feasibility, reliability and validity of such an approach are discussed.

## Materials and methods

### Participants

For this study, we recruited 19 neuro-typical adults aged from 18 to 26 years old (M = 22.3, SD = 1.9), among them 5 males.

### Procedure

Participants were welcomed into the lab and asked to sign a written consent form. The study was approved by the Ethics Committee of Psychology Research, University of Padua.

Participants sat on a desk and wore an accelerometer research watch on their dominant wrist (the experimental set-up is depicted in Figure 1). They were then asked to place the dominant hand at a specific starting position, monitored by a presence sensor. At the distance of their arm length, they found a response touchscreen so that they were required to completely extend their arm to touch the response screen. A specific task was proposed and required the participant to make action selection choices by touching one of the response keys on the screen. The task tested the participant’s ability to select a prepotent or an alternative response. During this behavioural task, the kinematics of participant’s dominant arm was monitored by the wrist worn 3-axis accelerometer. The task lasted about 15 minutes.

**Fig 1.**
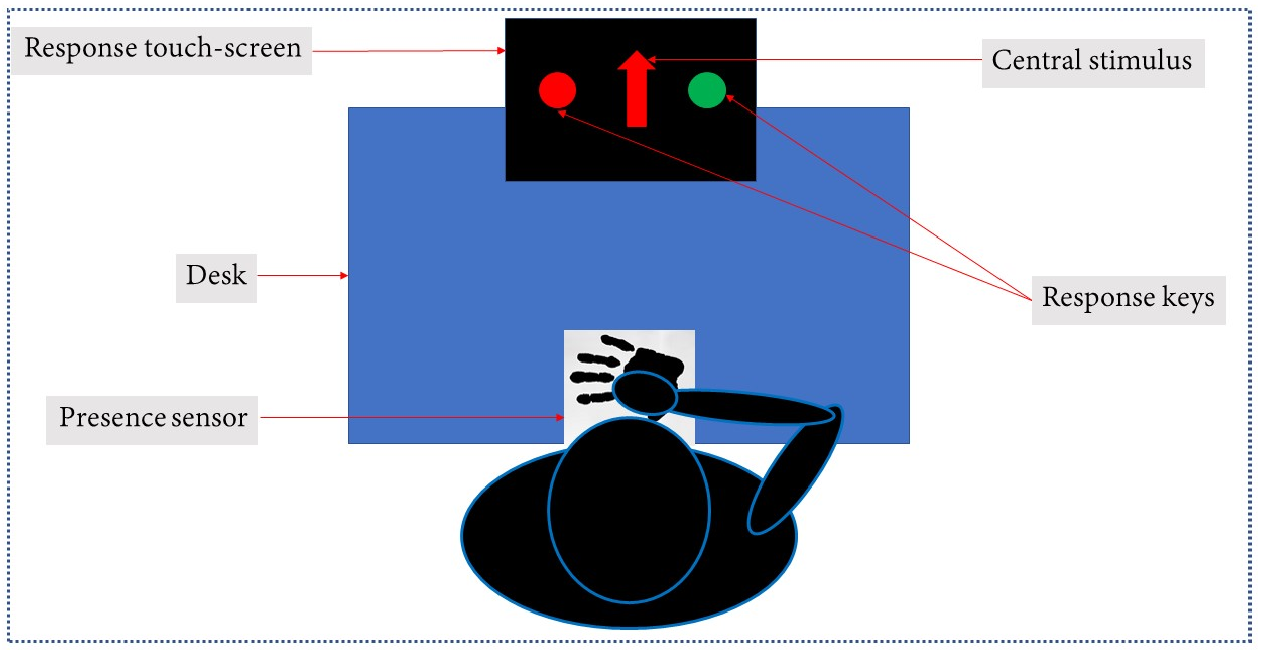
Experimental set-up.

### Apparatus

The dominant hand acceleration was monitored resorting to the GENEActiv Original 3-axis wrist worn accelerometer [20] (size: 43 mm × 40 mm × 13 mm, weight without the strap: 16 g). The device measured accelerations through a MEMS sensor, within a range of +/− 8 g, at a 12 bit (3.9 mg) resolution with a 100 Hz logging frequency.

The task was implemented resorting to a JavaFX based application [21].

To run the experiment, we employed a laptop Lenovo G50-80 (Intel Core i5-5200U (2.2 GHz), 4 GB DDR3L SDRAM, 500 GB HDD, 15.6” HD LED (1366 × 768), Intel HD Graphics 5500, Windows 10 64-bit).

The analysis of the resulting data was performed resorting to Python [22] and primarily to “pandas”, “numpy” and “scipy” libraries.

Participants responded by tapping on a 19 inch touchscreen (LG-T1910BP, response time 5 ms). The presence-absence of the participant’s hand on the starting position was detected through a custom-made presence sensor based on Arduino Leonardo which sent the hand detection data to the laptop via one of its USB ports. It was connected to a ground capacitor (100 pF) and a capacitive sensor, which consisted of a copper foil wrapped with plastic film (dimension 20 cm × 12 cm, thickness 0.1 mm). The presence sensor program was written using the Arduino Capacitive Sensing Library.

### Task: inhibition of prepotent response

A *Go/No-Go* paradigm was adapted to assess the inhibition of a prepotent response. In particular, on the arrival of a central stimulus (red/green, upwards/downwards arrow), participants were asked to select, reach and press one of two response keys (either a red circle or a green circle) placed one on the left and one on the right side of the central stimulus. Participants were told to select the response key of the same color of the central stimulus when it was an upwards/downwards (counterbalanced between participants) arrow (*dominant condition*). On the other side, they were told to select the response key of the different color when the central stimulus was an averted (either upwards or downwards, counterbalanced between participants) arrow (*non-dominant condition*). We built a prepotent response for the same-colour action, given that it was the one that appeared with a higher chance (75%). On the contrary, we elicited an inhibitory different-colour action, which was the less probable one (25%). In this way, we were able to measure the kinematics of dominant vs non-dominant selections, being the movements equal.

When participants failed to press any keys within 2,000 ms, the response was marked as “omission”. When participants moved their dominant hand from the starting position before the appearance of the cue stimulus, the response was tagged as “anticipation” and the program aborted the trial by providing no cue stimulus. For this task, each participant was required to perform 160 valid trials (i.e., trial with correct/incorrect answer). In any case, the total trials never exceed a maximum number of 180. Trials were divided into two blocks, distinguished by the red/green response keys being located once on the right and once on the left side of the touchscreen. To maintain participants’ attention and engagement during the task, a short video appeared every 40 trials.

Before the start of the next trial, the participant had to return his hand on the sensor. As soon as the hand was in place, as long as the previous trial was not running anymore, the next trial started after a random delay in the range from 0 to 2,000 ms. We will refer to this independent variable as StimulusRandomTime and analyse its effect on participants’ performance. Indeed, this variable manipulated the time available to pre-activate the sensorimotor system and predict the incoming occurrence of the central stimulus, potentially affecting the response timing [23].

## Kinematic analyses

For each trial with answer (i.e., no anticipation, no omission) we reported the time instants distinctive of the trial:

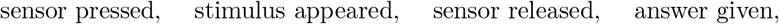

to whom we will from now on refer to as P, S, R, A.

To obtain these data, we combined the accelerometer data and the saved task outputs. To this aim, we synchronized the software logs and the accelerometer with the computer local time.

### Measures of interest

The time intervals that are related to the kinematic measures of interest were [*S, R*] which defined RT and [*R, A*] that corresponded to the MD and was used to compute the TPV%. In addition, the interval [*P, S*] determined the StimulusRandomTime.

We computed velocity from acceleration and then the Time to Peak Velocity percentage (TPV%), which is the percentage of time spent from R to maximum peak velocity with respect to the time interval from R to A (i.e., the MD). We employed the TPV% as a relative asymmetry index compared to the ideal symmetrical value of 50%, which would indicate an equivalent acceleration/deceleration phase. Based on the available literature, we expected the peak velocity to occur at around 40% of MD [24]. From the exploratory comparison between participants’ performance at the dominant and non-dominant conditions, a smaller TPV% will indicate a greater need for control and adjustment of the ongoing movement. On the other hand, a bigger TPV% will indicate a greater need for motor planning [1].

### Time to Peak Velocity detection

As described in detail in S1 Appendix, the effective acceleration is individuated by means of raw accelerometer data calibration and preprocessing. We are now interested to compute the TPV% value, in the following we walk through the adopted methodology.

From a theoretical and mathematical point of view, the most direct ways is to start to compute the TPV% applying an integration in time to obtain velocity from acceleration. In particular, let *a*(*t*) be the acceleration signal on one axis, the related velocity signal *v*(*t*) can be computed as follows:

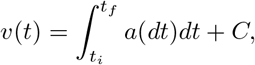

where *t_i_* and *t_f_* are the initial and final time instants of the movement and *C* is an integration constant.

However, when facing with real data and numerical functions (e.g., numerical integration), numerical errors can return unreliable velocity values.

Considering the calibrated and preprocessed acceleration, let *acc_RA_* be the signal related to the time interval [*R, A*] of a specific valid trial, we applied the cumulative trapezoidal numerical integration function in order to compute velocity. In Figure 2, we reported the velocity components obtained by applying this function to the acceleration ones of a trial. After this step, we also computed its magnitude (which represents the velocity module) from its components, also shown in Figure 2.

**Fig 2.**
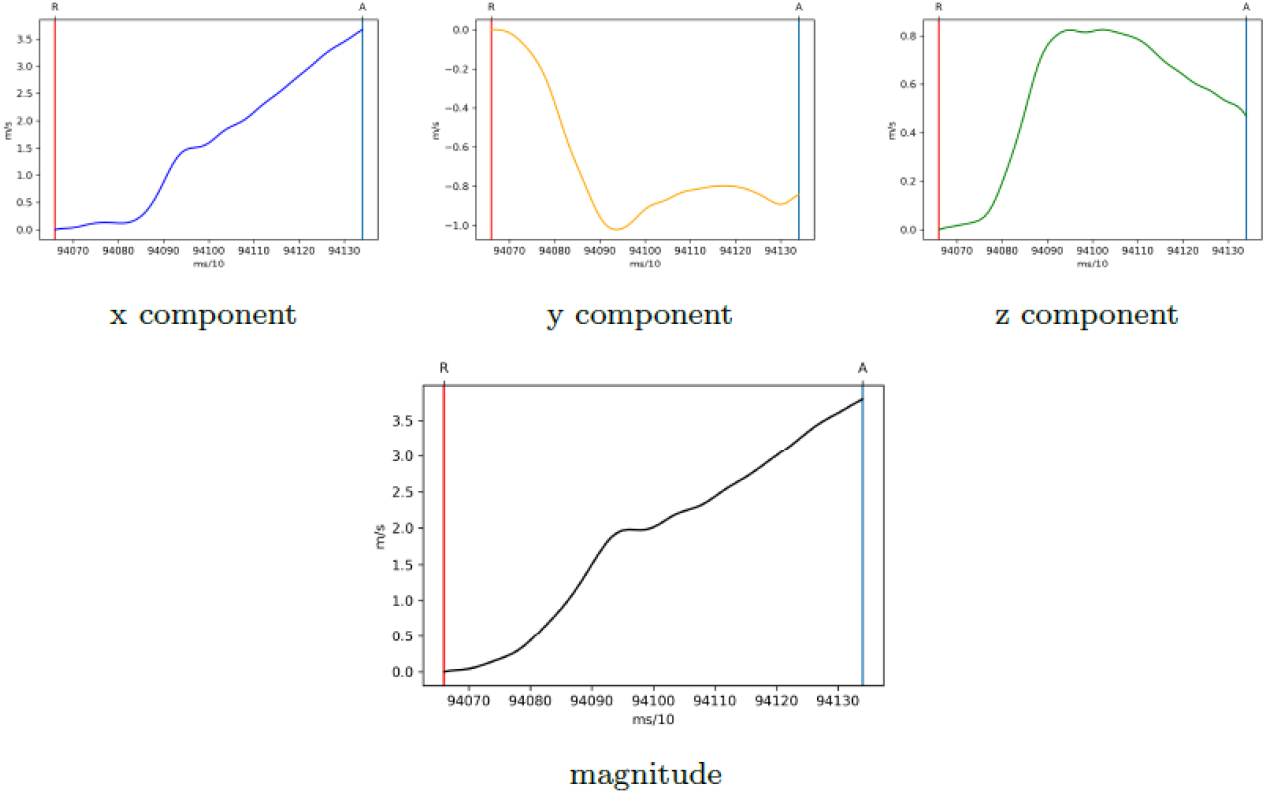
Velocity signals of a trial where the error due to the acceleration bias is visible in component x and in magnitude.

Notably, the application of an integration function could lead to an incremental numerical error due to a possible bias (i.e., additive noise) present in the acceleration, visible in Figure 2. Such phenomenon, may lead to the creation of a “new” and “false” maximum peak, making the computation of peak velocity quite challenging.

To overcome the aforementioned issue, we applied the detrend function to the velocity magnitude. The application of this function removed the signal linear trend and reduced the numerical error describe above (further details are reported in S2 Appendix).

In fact, while the velocity values could change due to the detrend function application, the position in time of the peak velocity appeared stable. Therefore, we decided to set aside the indices based on the velocity value (“how fast”, e.g., mean velocity, value of peak velocity) and to concentrate on the position in time of the peak velocity (“when” in time, e.g., time to peak velocity). This choice is supported also by the analysis of reliability and validity of the calibrated and preprocessed acceleration and the computed velocity values (S3 Appendix).

## Results

We were interested in exploring the differences between the *dominant* and *non-dominant* conditions, which stand for the difference between performing either a prepotent or an alternative response.

Participants performed 2,962 correct trials and 54 incorrect ones. We analysed the correct answers and computed the following indices of the participants’ performance at the inhibition task: Reaction Time (RT) from stimulus to movement onset (time interval [*S, R*]), Movement Duration (MD) from movement onset to answer (time interval [*R, A*]), Time to Peak Velocity percentage in the interval (0, 100) (TPV%) and the one rescaled within the interval (0, 1) (TPV).

We also aimed to exclude possible extreme TPV% values that would be due to numerical errors, in cases where the detrend function was not sufficient to remove their effect on the signal. Therefore, the a-priori inclusion criteria for valid TPV% values comprehended those between 5% and 95%. All values out of this range were considered invalid, namely not related to human movements but ascribable to technical failures. At the end of this procedure, we excluded 59 out of 2, 962 trials.

The following analyses have been run with R version 4.0.2 [25]. Means and Standard Deviations of RT, MD and TPV% are reported in Table 1.

**Table 1.** Means and Standard Deviations (*n_participants_* = 19).

| Condition | RT |  |  | MD |  |  | TPV% |  |  |
| --- | --- | --- | --- | --- | --- | --- | --- | --- | --- |
|  | n<sub>trials</sub> | M | SD | n<sub>trials</sub> | M | SD | n<sub>trials</sub> | M | SD |
| Dominant | 2,253 | 558 ms | 136 ms | 2,253 | 500 ms | 167 ms | 2,213 | 40% | 15% |
| Non-dominant | 709 | 601 ms | 163 ms | 709 | 591 ms | 207 ms | 690 | 45% | 17% |

*Note: TPV% includes less trials due to the exclusion of extreme values.*

From these descriptive statistics, we appreciate that participants performed the non-dominant condition (compared to the dominant one) by increasing the time devoted both to the motor planning (RT) and to the motor execution (MD).

In support of the validity of the TPV% index, our average values in the dominant (M = 40, SD = 15) and non-dominant (M = 45, SD = 17) conditions appeared consistent with those reported by previous studies and similar tasks [24]. Therefore, despite the high SD that questions the index reliability, we run further analyses to explore the factors affecting the TPV%.

To this end, a model comparison approach was adopted using the R package “glmmTMB” [26]. Mixed-effects models have been employed to account for the repeated measures design of the experiment (i.e., trials nested within participants). Generalized mixed-effects models were used considering the Beta distribution, with logit link function. Indeed, our dependent variable (TPV%) contained continuous proportions on the interval (0, 100), easily rescaled in the interval (0, 1) (TPV), and can be approximated by a Beta distribution [27]. The distribution of TPV values is shown in Figure 3.

**Fig 3.**
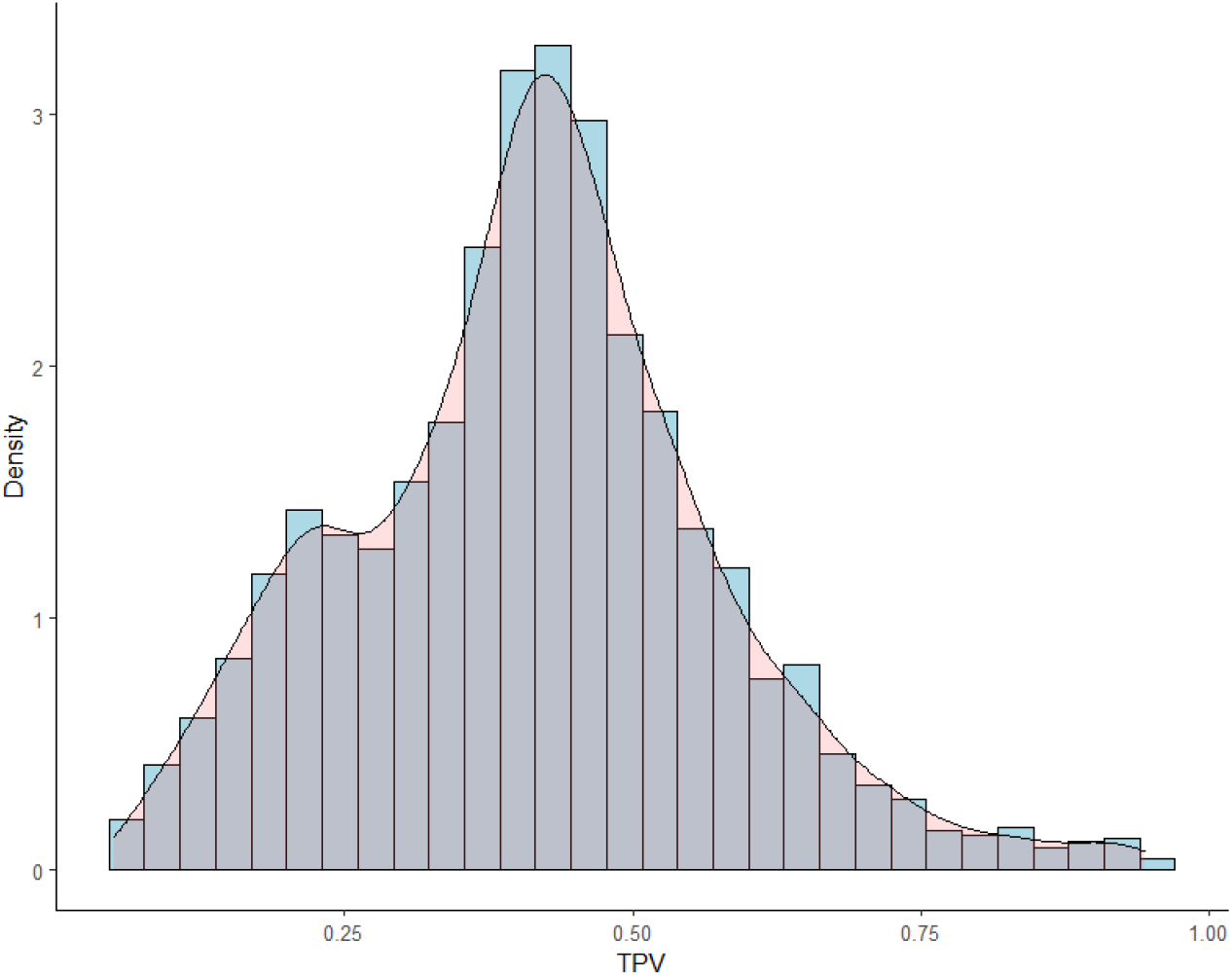
Distribution of the TPV values (*n_trials_* = 2, 903).

We aimed to account for the random effect of participants (i.e., interpersonal variability), as well as the fixed effect of condition (within-subjects, two levels categorical factor: dominant versus non-dominant). Moreover, we checked for the effect of the random time before the central stimulus onset. The latter was a continuous independent variable that we named StimulusRandomTime.

First of all, we specified four nested models to explore our data, with the TPV as dependent variable and the random effect of participants:

- `mb0` specified the hypothesis of no difference due to the independent variables and only accounted for the random effect of participants;
- `mb1` specified the hypothesis of a difference due to the condition effect;
- `mb2` specified the hypothesis of a difference due to the additive effect of condition and StimulusRandomTime;
- `mb3` specified the hypothesis of a difference due to the interaction effect of condition and StimulusRandomTime.

The details of the model specification are depicted in Table 2.

**Table 2.**
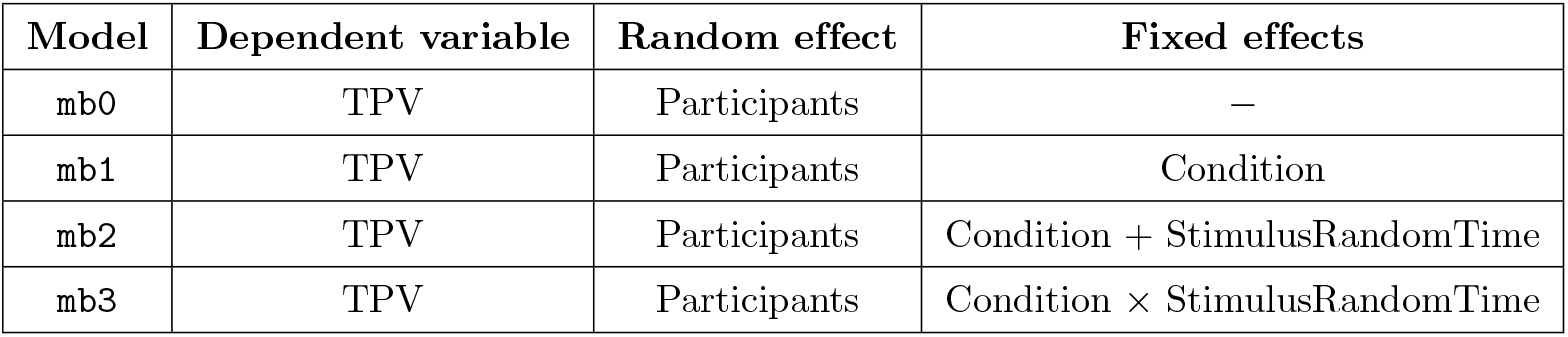
Model specification.

| Model | Dependent variable | Random effect | Fixed effects |
| --- | --- | --- | --- |
| mb0 | TPV | Participants | — |
| mb1 | TPV | Participants | Condition |
| mb2 | TPV | Participants | Condition + StimulusRandomTime |
| mb3 | TPV | Participants | Condition × StimulusRandomTime |

Therefore, the four models were compared using R package “AIC_cmodavg_” [28] to compute the Akaike weights (i.e., the probability of each model, given the data and the set of considered models) [29]. Moreover, the models have been compared using a likelihood ratio test (`anova(mb0, mb1, mb2, mb3)` R function). The degree of freedom (*Df*), the Akaike weights (*AICcWt*), the chi-squared test statistic values (*χ*^2^) and the p-values (*p*) are reported in Table 3.

**Table 3.** Model comparison.

| Model | $Df$ | $AICcWt$ | $\chi^2$ | $p$ |
| --- | --- | --- | --- | --- |
| mb0 | 3 | 0.00 |  |  |
| mb1 | 4 | 0.20 | 49.70 | <.001 |
| mb2 | 5 | 0.44 | 3.58 | .06 |
| mb3 | 6 | 0.36 | 1.63 | .20 |

The most plausible model given the data and the set of considered models was `mb2` (*AICcWt* = 0.44), which included the random effect of participants, the additive effects of condition (statistically significant according to *p* < .001) and StimulusRandomTime (statistically non significant according to *p* = .08) (p-values from `summary(mb2)` R function). These effects, predicted by model `mb2`, are depicted in Figure 4.

**Fig 4.**
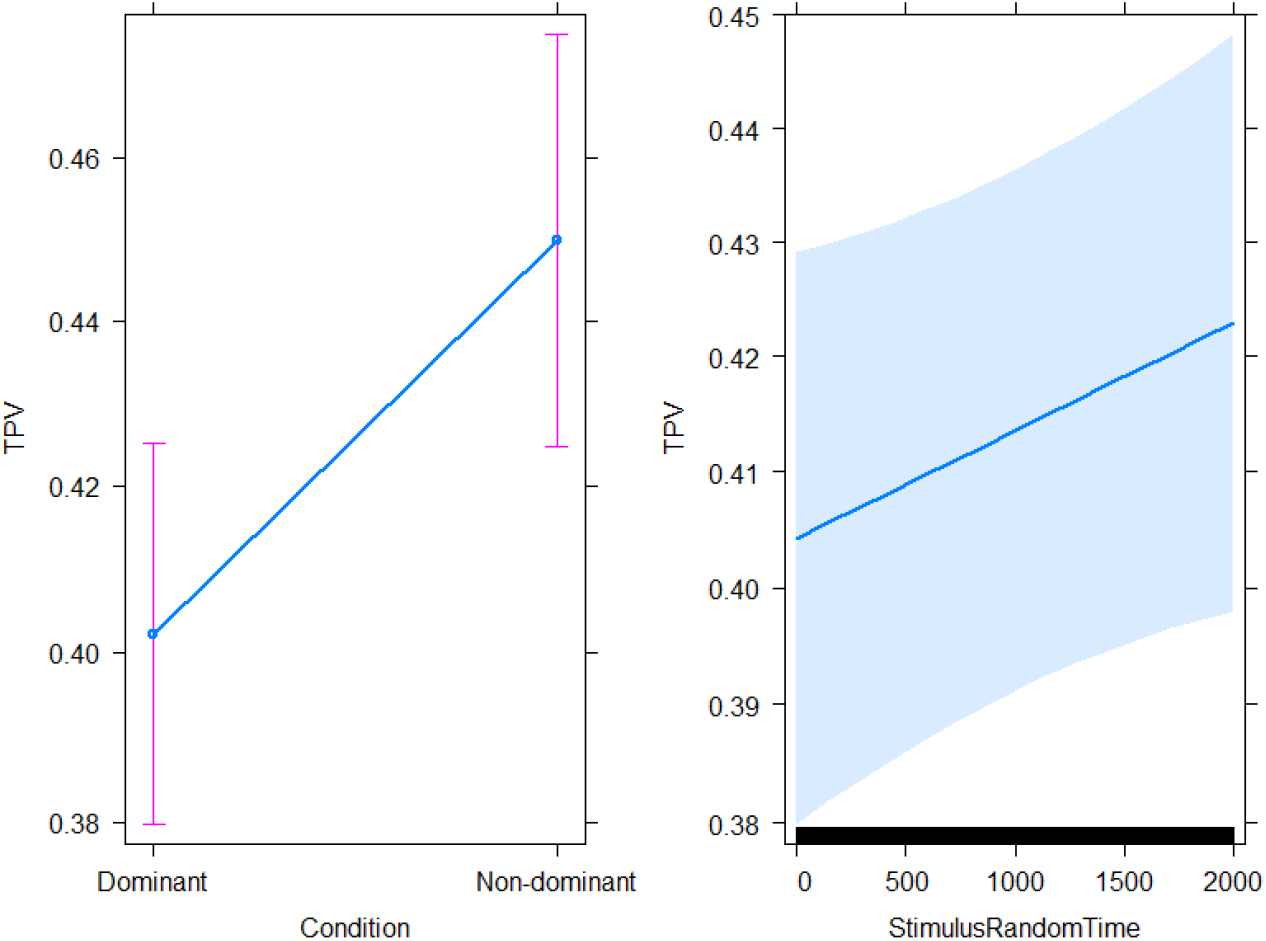
Model `mb2`: condition and StimulusRandomTime effects on the TPV (*n_participants_* = 19, *n_trials_* = 2, 903).

From the visualisation of the effects, we appreciate that the TPV was bigger (i.e., occurred later in time) in the non-dominant condition, with additional evidence that supports the higher need for motor planning rather than online adjustment. Moreover, the amount of time participants had to wait for the stimulus to appear (StimulusRandomTime) was positively correlated to the TPV. Although not significant from a statistical point of view, this effect suggests that a longer preparation time before the trial to start might allow participants to increase the time devoted to motor planning.

## Discussion

The present study explored the neurotypical adults’ movements in a task that required a reaching either to select a prepotent, dominant response or to inhibit the dominant and select the non-dominant alternative. The descriptive statistics indicated that participants performed the non-dominant condition (compared to the dominant one) by increasing both the RT (time devoted to the motor planning prior to movement onset) and MD (time of motor execution). However, these two indices are not sufficient to disentangle the planning and control phases of the movement. Indeed, given that motor planning and control overlap during the MD [1], we computed the TPV to further distinguish these two mechanisms. As a relative asymmetry index, whether the TPV occurred earlier or later over the MD would reflect more either planning or control. From our exploratory model comparison, we can expect people to have bigger TPV (which resulted in a shorter deceleration phase) in the non-dominant compared to the dominant condition. This evidence supported the idea that adults require a greater motor planning rather than online adjustment to inhibit a prepotent response, select and perform an alternative one.

In addition, the most plausible model given our data and set of specified models showed that when people had to wait more to start the trial, they increased the movement time devoted to motor planning. We can interpret this finding in light of the massive literature about the preparatory effect of the foreperiod, which is the time from a warning signal and a “Go” stimulus and affect response times [30]. In our study, participants had to place their hand on the presence sensor to signal their readiness to start the next trial. The time instant they pressed the sensor can be seen as an active warning signal that pre-activate the sensorimotor system. After a variable random time interval, the central “Go” stimulus appeared to trigger participants’ response. We can speculate that, within 2,000 ms, a longer preparation time increases adults’ motor planning. As the foreperiod effect and the temporal preparation abilities change across development, future studies could expand on the ontogensis of these mechanisms [31].

The present work also discussed the feasibility, potential and limits of a wearable 3-axis accelerometer to investigate human reaching movements. The inertial sensors built with 3-axis accelerometers, gyroscopes, and magnetometers have been indicated as promising commercial tools to study the kinematics of human movements and overcome the constraints of expensive motion capture systems. Although they have the potential of being portable and wearable, they appeared to provide accurate and reliable data only for some kinematic indices, such as the value and timing of peak velocity [10]. In light of these encouraging evidences, we employed a low-cost, easy to use, 3-axis wrist worn accelerometer to investigate the role of motor planning and control in a reaching to select or inhibit a prepotent response.

Based on our kinematic measurements and analyses, the kinematic indices built upon the velocity value did not appear sufficiently reliable and valid (as reported in S3 Appendix). On the other hand, those related to the velocity shape over time seemed to be valid indices. Indeed, our average Time to Peak Velocity percentage (TPV%) was consistent with those reported by previous studies, similar tasks and motion capture systems with highest level of precision [24]. Therefore, keeping in mind that the high standard deviation that questions the index reliability, we support the use of a commercial and low-cost 3-axis accelerometer to calculate the TPV% and compare within-participants’ performance.

From a methodological point of view, to further increase the accuracy of the preprocessing, in particular to remove the gravity component from the acquired acceleration, future studies could use a combination of accelerometer and gyroscope. In this way, data related to the orientation of the accelerometer would be available in order to remove the gravitational acceleration. However, the gyroscope would not solve the numerical errors driven by possible accelerometer bias and numerical mathematical functions. These issues could be addressed from an algorithmic point of view, with the evaluation of other methods and models in order to process raw accelerometer data in a way that could reduce the numerical errors. An algorithm class that could obtain promising results with huge amount of raw data is the learning class. Machine and deep learning algorithms could study different input signals and learning information from all the data. In this case, a supervised data set would incrementally improve the results but also an unsupervised approach could be taken into consideration.

Overall, this study expands on our understanding of which motor strategy is successful for neurotypical adults to perform or inhibit prepotent reaching movements. This would lay the foundations for investigating the atypical strategies implemented by individuals and clinical groups with inefficient inhibition. Moreover, the present approach could be adopted to investigate other mechanisms that intervene on action selection, motor planning and control. Further research is needed to develop and test theories examining the link between kinematics and neuropsychological mechanisms of action selection, which might be distinctive of several disorders and psychopathologies. Indeed, although difficulties and impairments in the domain of action execution are common to several clinical conditions, the underlying sensory, motor and cognitive mechanisms might dramatically differ among patients [32–35]. Future studies could utilise the present method and apparatus to disentangle the planning and control mechanisms of motor actions that involve different neuropsychological abilities.

## Supporting information

S1 Appendix. Acceleration calibration and preprocessing.

S2 Appendix. The detrend function application to velocity.

S3 Appendix. Reliability and validity of acceleration and velocity values.

## Acknowledgments

Our gratitude to the clever master students who collaborated to data collection: Sara Pezzotti, Ilaria Rossi, Irene Strappazzon. Many thanks to Andrea Janna for his fundamental technical support and advice.

