## Supplementary material for "Reaching to inhibit a prepotent response: a wearable 3-axis accelerometer kinematic analysis": S1 Appendix

### Acceleration calibration and preprocessing

The acceleration calibration and preprocessing analysis has been run on the data collected by an external experimenter (not part of the cohort involved in the trials) who repeated multiple selection tasks, just as a participant. Within each task, the experimenter answered to a central cue stimulus by tapping a central response key below the cue. In this way, the displacement remained roughly the same for each trial. In particular, the experimenter performed 40 trials: 1 anticipation, 2 omissions, 37 valid answers. The subsequent analysis focused on the raw acceleration signals that started when the sensor was pressed for the first trial and ended when the last valid answer was given.

The accelerometer data were sampled at 100 Hz (i.e., data sampled every 10 ms) and data were stored in g units for offline analyses.

Considering the 3-axis accelerometer, the principal output was, for each axis, the measured signal, which may be broken into the following components [1]:

$$\textit{acquired acceleration} = \textit{effective acceleration} + \textit{gravity acceleration} + \textit{noise}.$$

In order to examine the true movements of the participants, we processed the *acquired acceleration* components to obtain their corresponding *effective acceleration* ones, as raw acceleration signals also contained noise, which could include an offset error, and gravity. In particular, the separation of the latter components becomes increasingly difficult during rotational movements. In fact, in the case of rotational movements (which were observed during our experimental task), the frequency domains of the movement-related component and the gravitational component can overlap, thus their separation can become challenging [1].

Resorting to state of the art approaches [1], the effective acceleration was extracted implementing the following two key steps (Table 1): (a) a band-pass filter, and, (b) an offset estimation and subtraction step. We now proceed expanding the discussion regarding their use in this work.

| |
|---|
| raw accelerometer data |
| ↓ |
| band-pass filter |
| ↓ |
| offset estimation and subtraction |
| ↓ |
| conversion from g to m/s$^2$ |
| ↓ |
| calibrated and preprocessed acceleration |

**Table 1.** Acceleration calibration and preprocessing.

Following [1], a $4^{th}$ order Butterworth band-pass filter with cut-off frequencies equal to 0.2-15 Hz was applied to the signal. The filter cut-off frequency of 0.2 Hz was chosen on the presumption that most daily movements of human body parts occur at frequencies higher than 0.2 Hz. The cut-off frequency of 15 Hz was instead chosen to remove the effect of high-frequency noise. Also the 1-20 Hz cut-off frequencies were evaluated, considering other choices made in literature [1–4], however it was not possible to observe any meaningful difference with respect to the 0.2-15 Hz band. Comparing now the acceleration signals in Figure 1, it is possible to see that the raw acceleration components were shifted with respect to 0 g because of gravity. The z component, for example, would fall as low as −g. After applying the band-pass filter, all acceleration components adjusted to lie around 0 g.

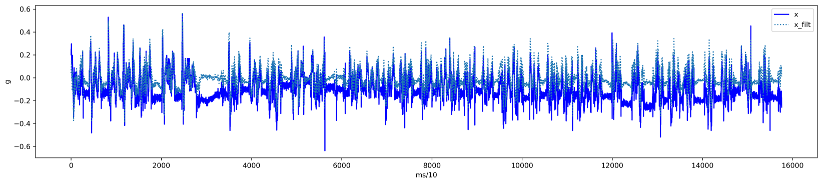


x component

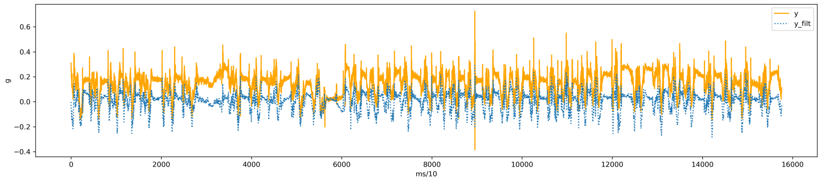


y component

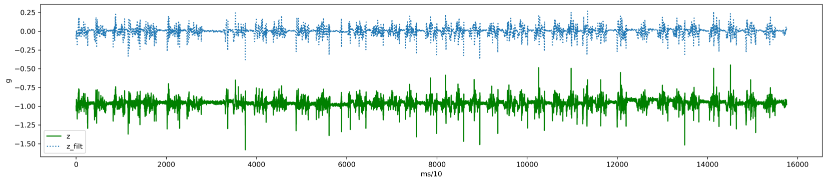


z component

**Fig 1.** Acceleration signals before (x, y, z) and after the band-pass filter (x_filt, y_filt, z_filt) application.

To estimate the offset error, data was collected from the accelerometer while at rest with the x, y and z axes pointing towards the ground (Figure 2).

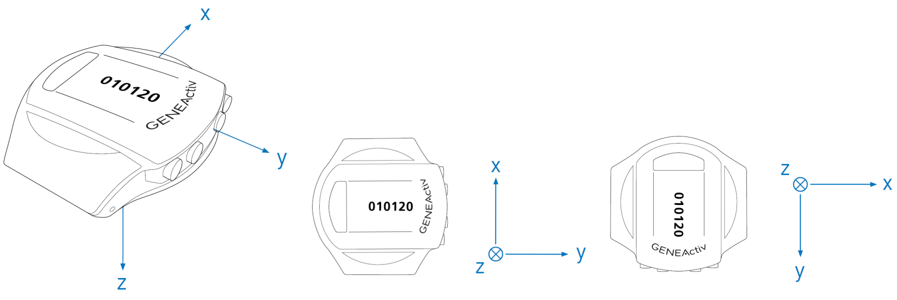


position 1 position 2 position 3

**Fig 2.** Accelerometer at rest positions [5].

From the filtered signal, for each of the three components, we computed the mean of the differences between actual accelerometer readings and the 0 g value expected from an accelerometer at rest. We hence obtained an offset value for each of the three axes. Successively, such values were removed from the acceleration data components, according to the pseudocode reported in Algorithm 1.

**Algorithm 1** Accelerometer offset

**procedure** (for each axis x, y, z)
  $i$ in $(x, y, z)$
  $df\_filt\_acc \leftarrow$ DataFrame with filtered acceleration
  $df\_offset\_acc \leftarrow$ new DataFrame for offset acceleration
  $epsilon_i \leftarrow$ offset value for axis i
  **for** $j$ in range (0, len($df\_filt\_acc$)): **do**
    **if** $df\_filt\_acc[j, acc_i] < (epsilon_i * (-1))$ **then**
      $df\_offset\_acc[j, acc_i] = df\_filt\_acc[j, acc_i] + epsilon_i$
    **else if** $df\_filt\_acc[j, acc_i] > epsilon_i$ **then**
      $df\_offset\_acc[j, acc_i] = df\_filt\_acc[j, acc_i] - epsilon_i$
    **else**
      $df\_offset\_acc[j, acc_i] = 0$

The visualisation of the signal from the accelerometer at rest fixed in the three different positions shows the filter effect and the presence of a offset error (Figure 3a and Figure 3b). Indeed, the offset removal led to data closer to zero (Figure 3c).

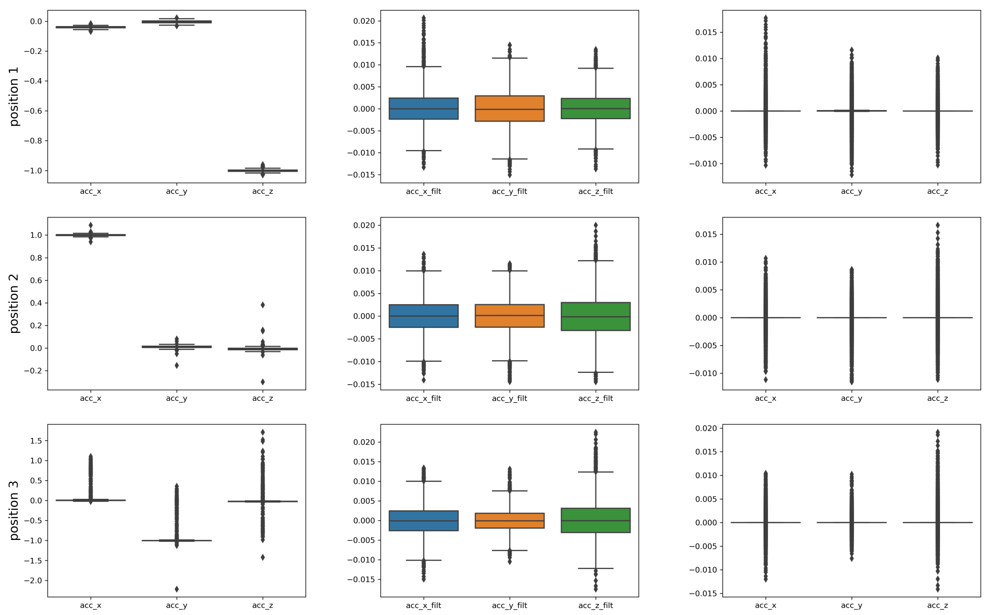


**(a)** **(b)** **(c)**

**Fig 3.** Acceleration values in g sampled at 100 Hz from the accelerometer at rest: (a) no filtering, (b) band-pass filtering, (c) band-pass filtering and offset removal ($n_{data}$ for each position $= 6,960$).

Finally, we obtained an estimate of the effective acceleration, adopting g = 9.80665 m/s$^2$ for the conversion from g to m/s$^2$ units [6].

5. GENEActiv. `https://www.activinsights.com/products/geneactiv/`.

6. Thorncroft, G. How precise is earth's gravity? `https://web.calpoly.edu/~gthorncr/ME302/documents/AccuracyofGravity.pdf`.
