## Supplementary material for "Reaching to inhibit a prepotent response: a wearable 3-axis accelerometer kinematic analysis": S2 Appendix

### The detrend function application to velocity

In the following analysis, with no loss of generality with respect to the aims of the procedure here described, we considered the exemplar waveforms $sin(t)$, $2 \cdot sin(t)$, $3 \cdot sin(t)$ as acceleration components signals. Therefore, we proceeded computing the velocity components integrating the acceleration ones and obtaining the velocity magnitudes reported in Figure 1a. After that, we applied the detrend function to the velocity magnitude, as shown in Figure 1b.

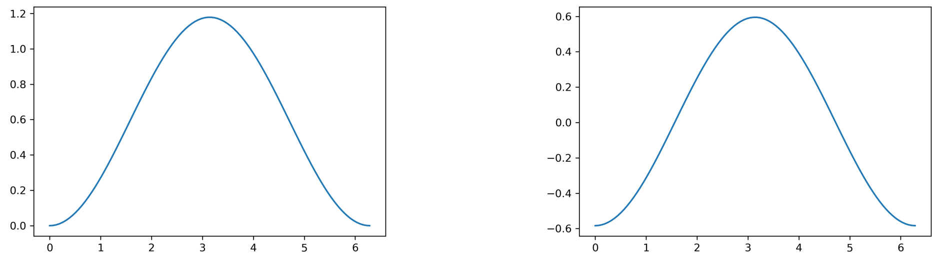


**(a)** **(b)**

**Fig 1.** Velocity magnitudes obtained from the integration of the acceleration vector components when no constant bias is present: (a) before applying the detrending and (b) after applying the detrending.

From these result it is possible to see that the application of detrend function only modified the signal respect to the ordinate axis but did not change the signal shape. This result is due to the fact that the velocity magnitude is computed from acceleration components characterized by neither trend nor bias.

Nevertheless, repeating the same analysis but starting from acceleration components, each of these with a constant bias, we obtained the velocity magnitude, before the application of detrend function, as shown in Figure 2a and, after the application of detrend function, as shown in Figure 2b.

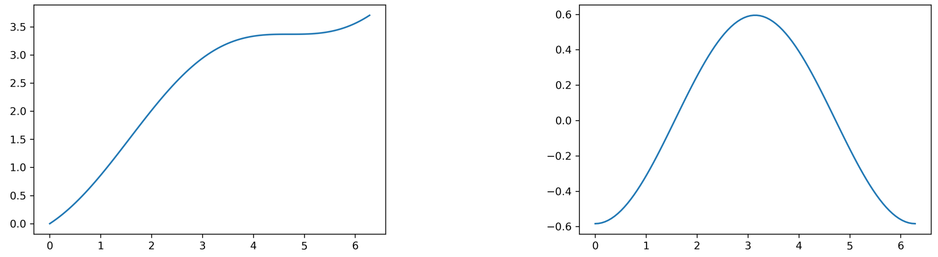


**(a)** **(b)**

**Fig 2.** Velocity magnitudes obtained from the integration of the acceleration vector components when a constant bias is present: (a) before the detrending and (b) after the detrending.

In Figure 2a it is possible to see an incremental numerical error due to the presence of the acceleration bias, as this is amplified by the application of the numerical integration function. Both the signal shape and the signal peak changed. Nevertheless, after the application of the detrend function, some of the signal changes due to this numerical error were removed, as reported in Figure 2b. In particular, it is important to note that comparing velocity signals in Figure 1b and Figure 2b: (i) the peak values changed, but (ii) the peak position in time is the same.

From the exploratory analyses on the signals, it is possible to draw the conclusion, hence, that although the velocity values could change due to the detrend function application, the position in time of the peak velocity remains stable. This property meets the requirement of individuating the TPV value set in this work.
