## Supplementary material for "Reaching to inhibit a prepotent response: a wearable 3-axis accelerometer kinematic analysis": S3 Appendix

### Reliability and validity of acceleration and velocity values

We finally worked to assess the reliability and validity of the calibrated and preprocessed acceleration and the computed velocity values.

As for the calibration and preprocessing analyses (S1 Appendix), we considered the data collected by an experimenter not belonging to cohort involved in our trials. We measured the distance between the sensor and where the response keys appear on the touchscreen, corresponding to the actual hand displacement required to reach the screen. We then compared such displacement to the one computed thanks to the wrist worn accelerometer.

In particular, we calculated the displacement of interest (i.e., from R to A) in three different ways. Under the hypothesis of constant acceleration, for each trial, we computed the mean acceleration from R to A and the displacement as the product between the mean acceleration and the square of time required to cover the distance of interest divided by 2 (i.e., according to the equation of uniformly accelerated motion). Under the hypothesis of constant velocity, for each trial, we calculated the mean velocity from R to A and computed the displacement as the product between time ($[R, A]$) and mean velocity (i.e., according to the equation of uniform motion). In addition, we computed the displacement by applying a double numerical integration to acceleration, using the cumulative trapezoidal numerical function, that does not rely on any hypothesis regarding acceleration or velocity.

It should be noted that to compute the displacement following the aforementioned procedures, the signal was not subject to detrending as in S2 Appendix. The detrending, in fact, can affect velocity component values, which is not acceptable when aiming to compute its magnitude. For this reason, the contribution of the numerical errors may be expected to appear in the displacement. The boxplot of the displacement values is visualised in Figure 1.

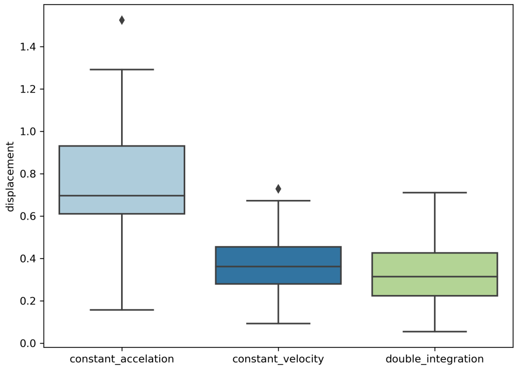


**Fig 1.** Displacement values in m computed from the acceleration values with different methods.

Then, we computed the mean and the standard deviation among all trials (37 trials with answer). For each method, these results are reported in Table 1 and compared to the actual displacement.

| Method | Mean | Standard Deviation | Measured Displacement |
|---|---|---|---|
| Constant acceleration | 0.76 m | 0.28 m | 0.46 m |
| Constant velocity | 0.38 m | 0.15 m | |
| Double integration | 0.34 m | 0.16 m | |

**Table 1.** Computed and actual displacement values ($n_{trials} = 37$).

Notably, the mean values are distant from the actual displacement and the standard deviations are quite high, especially under the hypothesis of constant acceleration. This could be due to the fact that the assumption of neither a constant acceleration nor a constant velocity are really appropriate to the actual characteristics of our task. Moreover, a double integration to compute displacements from acceleration can lead to large numerical errors, making this a weak method to assess the reliability and validity of velocity values. Indeed, this computation could be principally impeded by the accumulation of the numerical errors discussed so far.

Concluding, with this work we were not able to confirm nor disprove the reliability and validity of acceleration and velocity values obtained from a setting based on a wrist worn sensor. Nevertheless, we were able to show that such approach may be put to good use to instead obtain the peak velocity timing ("when" in time, e.g., time to peak velocity), information that may be fruitfully used to analyze the response of a subject.

### Supporting information

**S1 Appendix. Acceleration calibration and preprocessing.**

**S2 Appendix. The detrend function application to velocity.**
